# Significantly amplified photoacoustic effect for silica-coated gold nanoparticles by interface heat transfer mechanisms

**DOI:** 10.1101/2022.09.28.509922

**Authors:** Jonghae Youn, Peiyuan Kang, Blake A. Wilson, Chen Xie, Lokesh Basavarajappa, Qingxiao Wang, Moon Kim, Kenneth Hoyt, Zhenpeng Qin

## Abstract

Plasmonic gold nanoparticles (AuNPs) are effective photoacoustic (PA) signal agents and have found important biomedical applications. The silica coating on the surface of AuNPs showed enhanced PA efficiency, however, the PA amplification mechanism remains unclear. Here, we systematically studied the silica coating effect on PA generation of AuNPs under different laser pulse durations. We experimentally demonstrated up to 4-fold PA amplification under thin silica coating (<5 nm) and a picosecond laser excitation. The theoretical model further suggests that the PA amplification originates from two interface heat transfer mechanisms including 1) the enhanced interface thermal conductance on the silica-water interface and 2) the electron-phonon energy transfer channel on the gold/silica interface. This study discovers a regime of large PA amplification and provides a new rationale for plasmonic nanoparticle design to achieve better PA efficiency.

## INTRODUCTION

The photoacoustic (PA) effect is the generation of sound waves following light absorption and heating in a material. Upon light pulse irradiation, the material converts light energy into thermal energy and subsequently causes thermoelastic expansion of the material and the surrounding medium.^1,2^ PA effect has found wide applications including in the biomedical field since it reveals anatomical, molecular, functional, metabolic, and mechanical information of the cells and tissue.^3–8^ Exogenous PA contrast agents are often utilized to provide information on cellular and molecular events.^9–12^ To achieve an accurate diagnosis, the high optical-to-acoustic conversion efficiency is crucial for exogenous PA contrast agents.^13–17^Among diverse PA contrast agents, plasmonic nanoparticles have been widely used due to their strong optical absorption, lack of photobleaching, and easy surface functionalization.^13,18,19^ Further engineering of the nanoparticle surface can improve signal generation efficiency and sensitivity, ^24–29^ potentially expanding the applications of plasmonic nanoparticles in PA imaging. For example, surface coatings such as polydopamine, melanin, and graphene oxide can increase the nanoparticle absorbance and thereby increase PA signal amplitude,^20–22^ while coatings with large thermal expansion coefficients like PDMS can also enhance nanoparticle PA signal generation.^14^

For plasmonic gold nanoparticles (AuNPs), coating the particle surface with a shell of silica has been shown to yield up to 3-fold amplification of the PA signal.^30-34^ Enhanced interface heat transfer across the particle-water interface has been proposed as one potential explanation for the PA enhancement of silica-coated AuNPs.^30-32^ A study by Hu et al. reported time-resolved spectroscopy measurements of spherical AuNPs after femtosecond laser pulse stimulation that indicate thin silica coatings (<=10 nm) increased the heat dissipation rate from 15 nm spherical AuNPs into water, compared with bare AuNP. ^24^ This study suggests that the silica layer enhanced the interface heat transfer across the particle-solvent interface. However, theoretical modeling work from Shahbazi et al.,^34^ as well as Pang et al.,^35^ suggests that the limited heat conduction of the silica layer would lead to decreased PA signal for spherical AuNPs under nanosecond (ns) laser pulses. Recent theoretical modeling work suggests that an additional heat transfer channel from electron-phonon (e-ph) coupling across the gold-silica interface of spherical silica-coated AuNPs could facilitate enhanced interface heat transfer and lead to amplified PA signal for particles with a thin shell (< 10 nm) of silica under ps laser pulses at low fluence (i.e., below the threshold for cavitation effects).^26,43^ Regardless, with conflicting experimental results on whether silica coating amplifies or reduces the PA signal of spherical AuNPs stimulated by nanosecond (ns) laser pulses,^32,25^ and a lack of studies investigating the PA response of silica-coated AuNPs under ps laser pulse stimulation, it’s unclear what mechanism best describes the PA signal generation of spherical silica-coated AuNPs.

In this paper, we report up to 4-fold PA amplification for spherical AuNPs coated with a thin shell of silica (< 5 nm) under a picosecond (28 ps) laser excitation. We demonstrate that this amplification is highly sensitive to silica coating thickness and the laser pulse duration. The PA enhancement by silica coating originates from two interface heat transfer enhancements upon silica coating: 1) the enhanced interface heat conductance between silica and water and 2) the direct energy transport channel between gold electron and silica phonon at the gold-silica interface. Our work reports a new regime of large PA amplification under ps laser excitation and elucidates the possible mechanism for the PA amplification of metal-dielectric core-shell nanoparticles.

## Results

First, we prepared monodisperse silica-coated AuNP (Au@SiO_2_) and a flow system for stable PA measurements. Here we followed a method reported by Liz-Marzan’s group^36^ that utilized sodium silicate as pre-coating (Figure S1) to minimize the introduction of additional substances other than silica. Spherical AuNPs and uniform surface coating of silica on AuNPs were confirmed by transmission electron microscopy (TEM) with uniform 4 nm silica coating (Figure 1A). We further confirmed the uniform distribution of Au@SiO_2_ by dynamic light scattering (DLS, Figure 1B) to minimize the PA signal from nanoparticle aggregation.^37^ The plasmon resonance peak of Au@SiO_2_ shows a 4 nm red shift due to the refractive index change by silica coating (Figure 1C). To avoid the impact of AuNP modification and possible loss of PA signal with multiple laser pulse irradiation,^38^ we designed a flow system that allows PA signal is generated from fresh AuNPs (Figure 1D). The flow rate was determined by letting the PA signal reach a plateau (Figure S2) and particles are irradiated less than 4 pulses on average. With these conditions, we assessed the effects of laser duration and silica coating on the PA signal generation. We compared the PA signals from 15 nm bare AuNP and Au@SiO_2_ with 4 nm silica shell under ps and ns laser excitations (Figure 1E, F). Upon the ps laser irradiation, the signal amplitude of Au@SiO_2_ is larger than that of bare AuNP. However, no noticeable PA signal improvement was observed from ns laser. We further analyzed the peak-to-peak (P2P) amplitude of PA signal with different laser fluence for ns and ps lasers. The linear curves suggest the linear PA signal regime for both ps and ns laser excitations. With the ps laser excitation, the slope of P2P amplitude for Au@SiO_2_ is 4.3-fold higher than that for the bare AuNP (Figure 1G), while there is no significant enhancement from the ns laser (Figure 1H). The strong PA signal amplification is also observed in a larger 45 nm Au core with silica coating under ps laser with a 2.8-fold higher slope (Figure S3).

**Figure 1.**
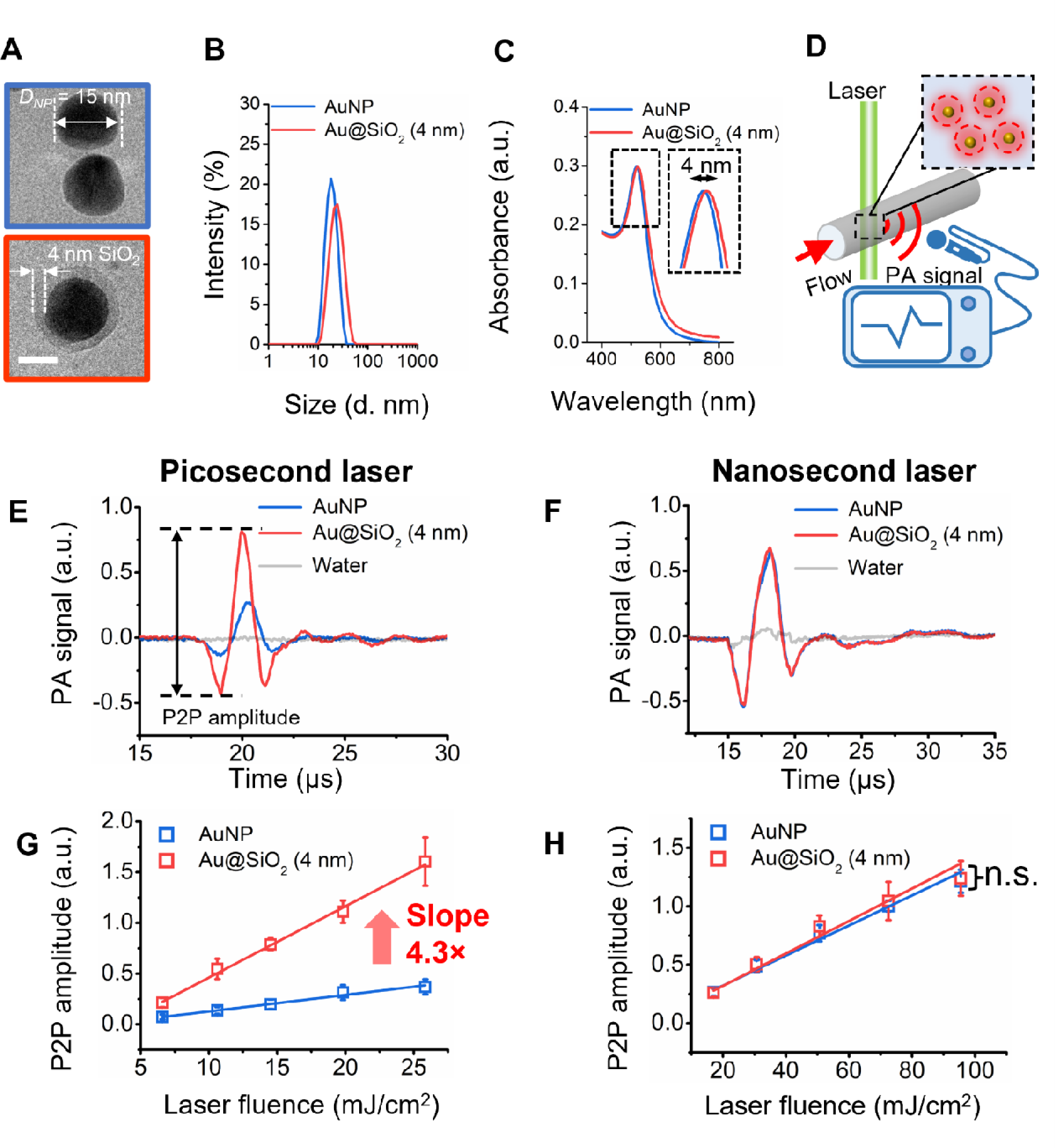
Photoacoustic signal amplification from silica-coated gold nanoparticles. (A) TEM images of 15 nm bare AuNP (blue box) and the AuNP with 4 nm silica coating (Au@SiO_2_, red box). Hydrodynamic size distribution (B) and UV/Vis spectra (C) of bare AuNP and Au@SiO_2_. (D) Schematic illustration of the flow system for photoacoustic signal amplification. Photoacoustic wavefunctions of 15 nm bare AuNPs and Au@SiO_2_ induced by 19.7 mJ/cm^2^ picosecond laser (E) and 72.5 mJ/cm^2^ nanosecond laser (F). The amplitude of photoacoustic signals of 15 nm bare AuNPs and Au@SiO_2_ induced by different laser fluences of picosecond laser (G) and nanosecond laser (H). n=30 measurements; the error bars show the standard deviation.

Second, we systemically studied the effect of the silica shell thickness on the PA signal generation with 15 nm and 45 nm Au cores under ps and ns laser. Specifically, we prepared silica coatings from 2 nm to 16 nm thickness. It remains challenging to synthesize monodisperse Au@SiO_2_ with silica coating thinner than 2 nm or thicker than 20 nm due to problems of particle stability and purity (Figure S4). TEM imaging shows uniform silica coating thicknesses for 15 nm Au core (Figure 2A) and 45 nm Au core nanoparticles (Figure 2C), respectively. The plasmon resonance peak shows a small red shift after silica coating (Figure S5 and Table S1), consistent with a previous report.^39^ For the 15 nm Au core (Figure 2B), the first 3 thin coatings (2.5 ± 0.5 nm, 3.4 ± 0.7, and 3.9 ± 0.7) show large PA enhancement (Eq. 1, SI) compared to that of bare AuNP, but no enhancement was observed in thicker silica coatings. The enhancement for the 45 nm Au core was observed at coating thicknesses of 2.0 ± 0.4 nm, 3.6 ± 0.6 nm, 3.8± 0.8 nm, and 4.9 ± 1.2 nm (Figure 2D). Among these, the largest amplification we observed was ∼400% enhancement from the 15 nm Au core with 4.6 nm silica coating and ∼300% enhancement from the 45 nm Au core with 4.9 nm silica coating. Overall, the highest amplification was observed in thin coating thickness (< 5 nm) as the amplification diminishes for thicker coating thicknesses (> 5 nm).

**Figure 2.**
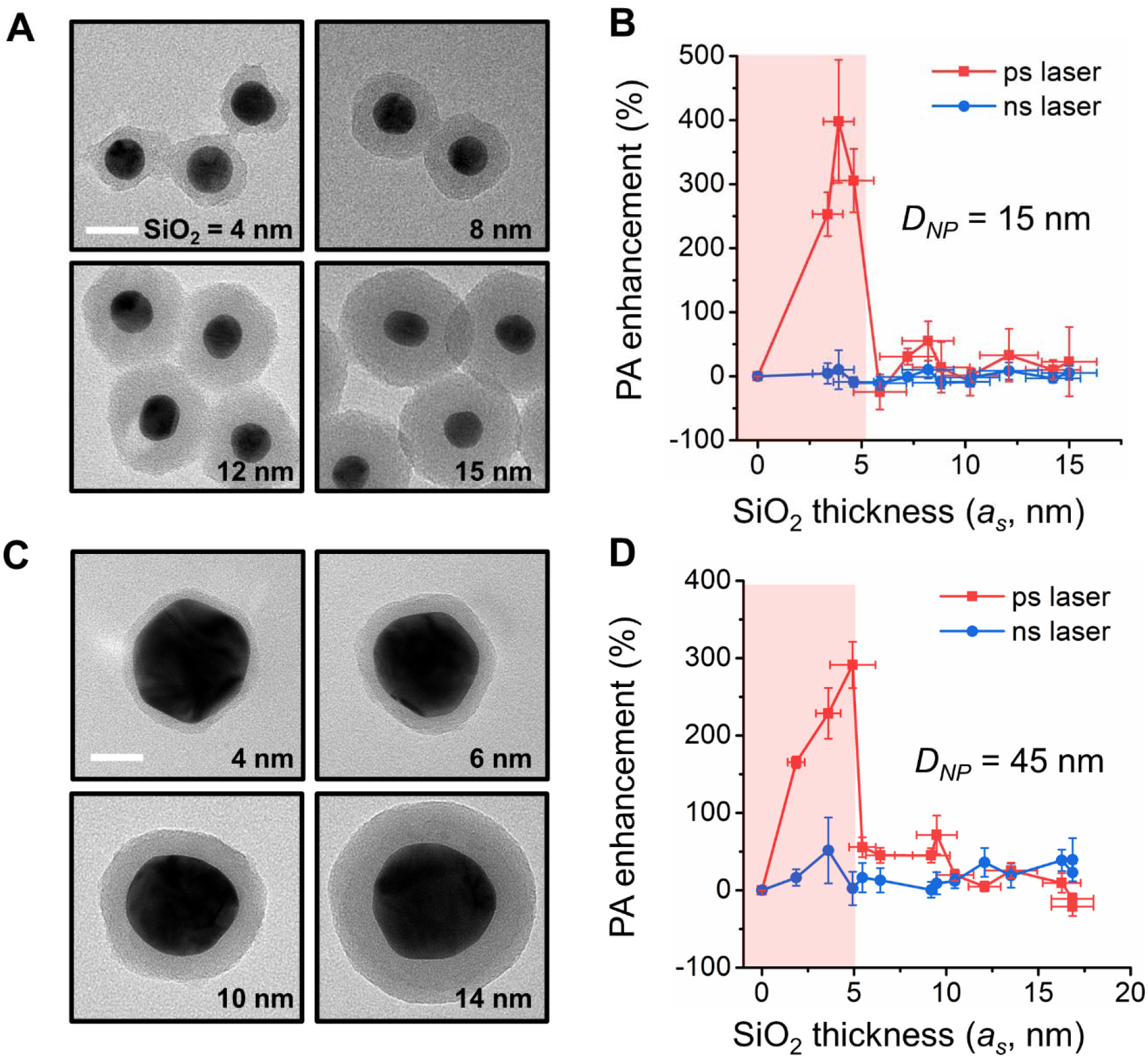
Photoacoustic signal amplification is limited to thin SiO_2_ coating thicknesses. TEM images of (A) 15 nm core and (C) 45 nm core gold nanoparticles with different coating thicknesses. Scale bar, 20 nm. Effects of SiO_2_ coating thickness and laser duration (ps laser: 19.8 mJ/cm^2^, ns laser: 72.5 mJ/cm^2^) on PA signal generation from (B) 15 nm Au core and (D) 45 nm Au core. All data points were merged from 3 different experiments (n=3). The red shades indicate the range of silica shell thickness with PA amplification.

Next, we investigated the effect of electron-phonon coupling on PA amplification. Optical energy is typically transported through electrons to lattice phonons within tens of ps. When the laser duration is on the order of ps, the nonequilibrium between electrons and phonons is significant and the electron temperature is much higher than the lattice temperature. As such, electron-phonon (*e-ph*) coupling plays an important role in the nanoparticle heating profile during ps laser excitation. ^34,35,42^ In contrast, when the pulse duration is on the order of ns *e-ph* coupling between the electrons and lattice can typically be neglected because the electron and phonon temperatures tend to remain at quasi-equilibrium during the heating process.^34,35,40^ A recent report by Alkurdi *et al*. suggests that the energy transfer by *e-ph* coupling on the interface of gold/silica can significantly alter the heating dynamics of both silica-coated gold nanoparticles and the surrounding medium.^26^ In this case, electron energy can be transported to the silica shell through two channels. The first channel is through gold phonons to silica phonons by the phonon-phonon (*ph-ph*) interaction at the gold-silica interface. The second channel allows energy to be directly deposited to silica phonons through the electron-phonon (*e-ph*) coupling on the gold-silica interface (Figure 3A). However, the effect of the additional *e-ph* heat transfer channel on PA signal generation under ns pulse laser is negligible below the threshold fluence for cavitation,^43^ so it can also be neglected for ns laser excitation. For the bare AuNP, the energy is only transported from gold phonon to water phonons through the phonon-phonon interaction at the AuNP-water interface (Figure 3B). For ns laser heating simulations, we thus used a single-temperature model without *e-ph* coupling that is like the model used by Pang et al.^35^ For ps laser heating, we adapted a previous hyperbolic two-temperature model and extended it to include silica coating with *e-ph* coupling on the interface of gold/silica like that reported by Alkurdi *et al*.^41,26^ The effect of silica coating on the heating process is demonstrated by the temperature distribution of Au@SiO_2_ with a 15 nm AuNP core (*D*_*NP*_ = 15 nm) and the direct *e-ph* channel at the interface (Figure 3C). By considering the direct *e-ph* heat transfer, the heating rate of surrounding water was significantly enhanced compared to the case with *ph-ph* channel only (*R*_*es*_ =∞) and the case with the bare nanoparticle (Figure 3D and S6A). Also, the heating rate of water decreases with silica shell thickness due to the heat absorption of the silica shell (Figure S6A). By considering the *e-ph* channel at the interface of gold/silica, the PA waveform of Au@SiO_2_ (a_s_ = 4 nm) has a higher amplitude than that of bare AuNP. In other words, the 4 nm silica coating can amplify the PA generation, consistent with our experimental observation. Furthermore, the PA enhancement by silica coating is sensitive to silica shell thickness (Figure 3E and S6B). The PA quantum yield (*Ф*_*PA*_) was defined as the ratio between the emitted PA energy and input optical energy (Eq 11, SI).^42^ We found that 2 nm coating of silica can lead to a 6-fold enhancement of *Ф*_*PA*_, compared to bare AuNP (Figure 3F). The *Ф*_*PA*_ enhancement decreases with silica thickness due to the heat absorption of silica coating, with no PA amplification for coating thickness larger than 5.7 nm. This thickness-dependent PA amplification result agrees with our experimental findings (Figures 1 and 2). We further estimated the PA generation under the ns laser. Interestingly, the PA quantum yield of Au@SiO_2_ particles under ns laser remains constant compared with that of bare AuNP. This is expected because the energy transport through the *e-ph* channel on the interface is primarily dependent on the temperature differences between electron and phonon systems, which is only significant at the time scale comparable to the *e-ph* interaction time (several ps). These modeling results demonstrated that the enhancement of the heating rate in the medium requires a thin silica coating and leads to a significantly higher PA generation efficiency during ps laser excitation.

**Figure 3.**
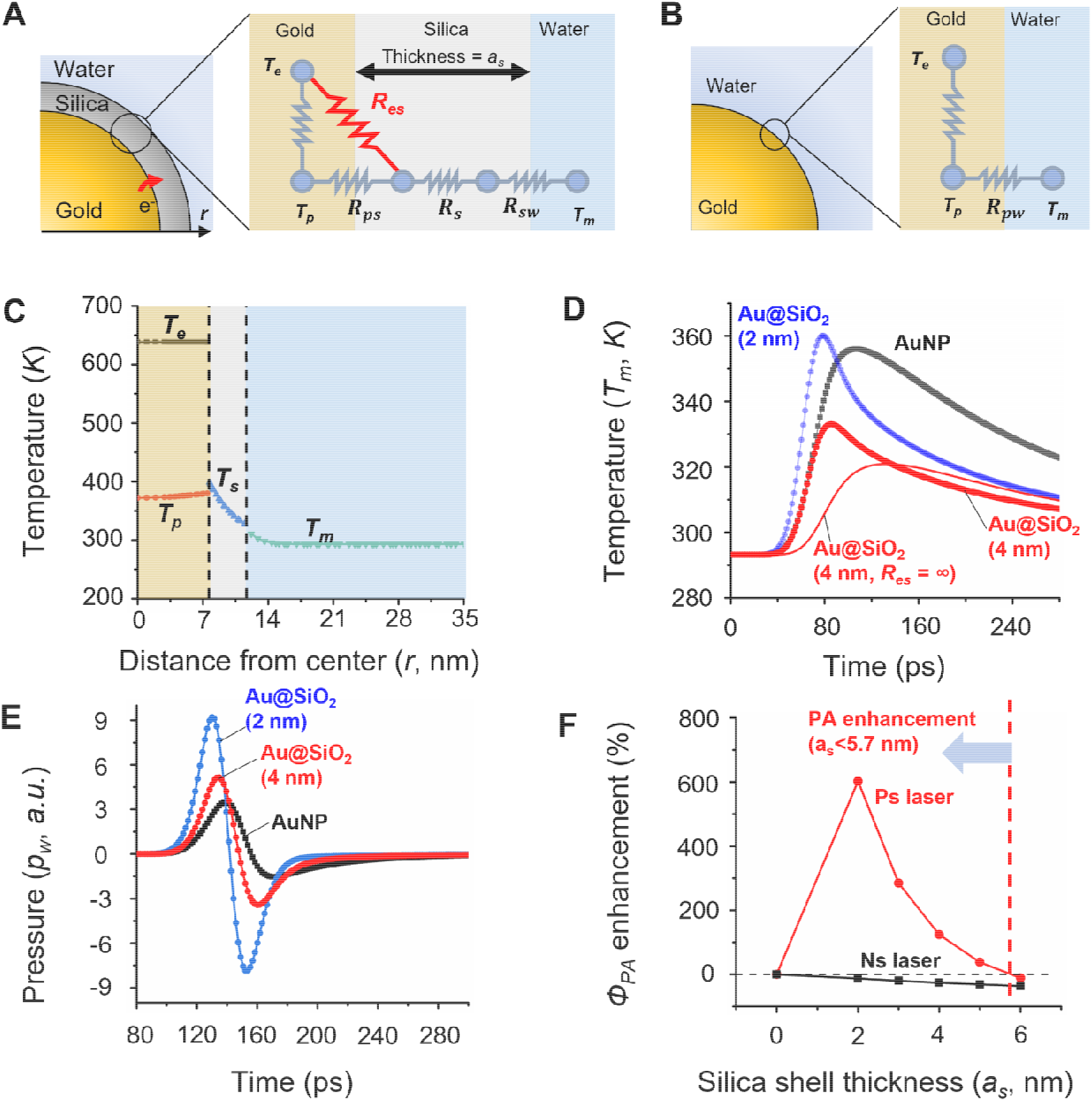
Gold-silica electron-phonon coupling contributes to photoacoustic amplification at thin SiO_2_ coating thicknesses. (A-B) Heat transfer model for (A) silica-coated gold nanoparticle (Au@SiO_2_) and (B) bare gold nanosphere (AuNP). Compared with the gold/water interface, an energy transport channel is included from gold electron to silica phonon at the interface of gold and silica (*R*_*es*_). (C) The temperature profile for Au@SiO_2_ (Au core is 15 nm, and silica thickness a_s_ = 4 nm) at 70 ns time point. The full width half maximum (FWHM) of the laser pulse is 28 ps. (D) The temperature evolution for water at the surface of Au@SiO_2_ and the bare AuNP. The “*R*_*es*_ = ∞” indicates the case only considering the phonon-phonon transport channel on the gold/silica interface. (E) PA wavefunctions of Au@SiO_2_ (a_s_ = 4 nm) and AuNP with the electron effect considered. (F) The silica shell thickness effect for PA quantum yield efficiency (*Ф*_*PA*_) under ps and ns laser pulses.

Lastly, we evaluated the impact of interface conductance on PA amplification. The heat transfer across two materials will lead to a temperature discontinuity at the interface that is characterized by interface thermal conductance (ITC, *h*). The silica-water ITC (*h*_*sw*_, 150 MW m^-2^ K^-1^)^43^ has been reported to be on the same level as ones for the gold-water interface (*h*_*pw*_, 105 MW m^-2^K^-1^)^44^ and gold-silica interface (*h*_*ps*_, 141 MW m^-2^K^-1^).^26^ However, a recent study suggested that the silica-water interface has thermal conductance more than 10 times higher (*h*_*sw*_, 2300 MW m^-2^K^-1^) than that of gold-water or gold-silica interfaces due to favorable hydrogen bonding effects.^45^ Hydrogen bond enhanced interface conductance has also been reported for simulations of hydrophilic, hydroxyl-terminated, self-assembled monolayers.^46^ To simplify the comparison, we defined the equivalent thermal resistance (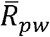, Eq 13 – 15, SI) for Au@SiO_2_ that incorporates the gold-silica and silica-water interface conductance, as well as the thermal resistance of the silica shell (Figure 4A). Here we keep consistent with our experimental conditions using thin silica coating thickness (*a*_*s*_ = 4 nm) and citrate-coated 15 nm AuNP (*R*_*pw*_ = 13.5 MK/W).^44^ Since the coating thickness and thermal conductivity for silica are fixed, the interface resistances for gold/silica (*R*_*ps*_) and silica/water (*R*_*sw*_) determine interface heat transfer and PA signal. To provide an overview of how the gold/silica and silica/water interfaces determine the overall heat transfer compared with bare AuNP, Figure 4B shows the combination of gold/silica and silica/water ITC that give an equivalent overall thermal resistance with bare AuNP 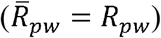.

**Figure 4.**
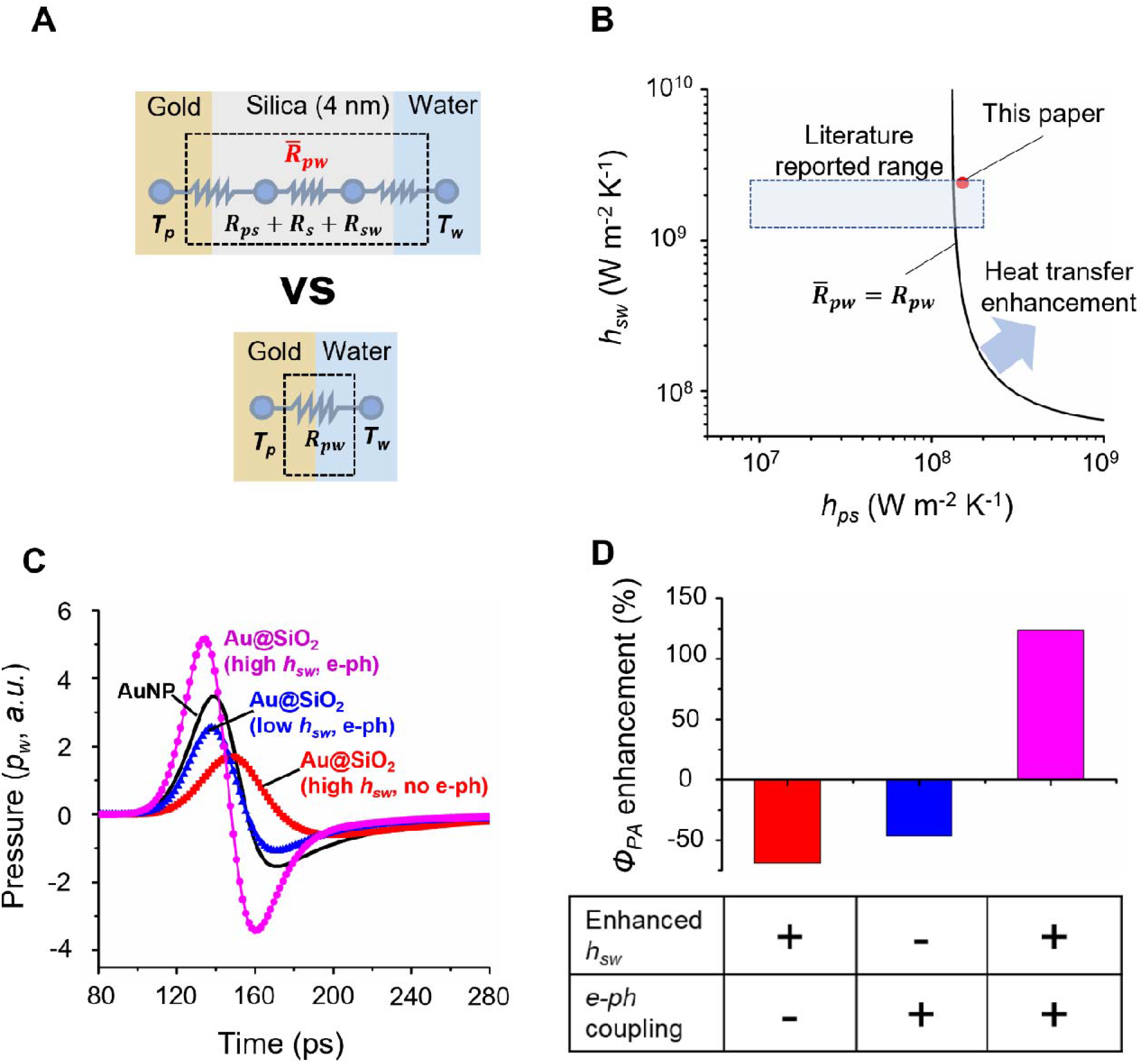
The PA enhancement requires the synergy of enhanced silica-water interface thermal conductance and gold-silica electro-phonon coupling. (A) The schematic of equivalent thermal resistance () for silica-coated AuNP (Au@SiO_2_). R and T represent thermal resistance and temperature, respectively. Subscripts “ps”, “sw”, and “pw” indicate the interface of gold/silica, silica/water, gold/water, respectively. The Subscript “s” indicates the silica domain. The equivalent thermal resistance of whole silica coating is represented as. (B) The interface thermal conductance on gold/silica (*h*_*ps*_) and silica/water (*h*_*sw*_) interface for an Au@SiO_2_ (silica thickness is 4 nm). The solid line represents the combination of *h*_*ps*_ and *h*_*sw*_ to allow 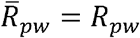. The shaded area is the range for *h*_*ps*_ and *h*_*sw*_ in previous reports.^45,46^ The red dot indicates the interface thermal conductance of the silica coating used in this study. (C) The pressure waveform of AuNP and Au@SiO_2_ (4 nm) under ps laser with considering with either enhanced *h*_*sw*_ or *e-ph* coupling effect. (D) Synergistic heat transfer enhancements on gold-silica interface (*e-ph* coupling) and silica-water interface (enhanced *h*_*sw*_) leads to the PA enhancement observed in the experiment.

Regions above the solid line show an enhanced heat transfer, and the shadowed area illustrates the reported range of conductance values.^45^ The combination of gold-silica and silica-water ITCs used in our model (*h*_*ps*_ =141 MW m^-2^K^-1^, *h*_*sw*_=2300 MW m^-2^K^-1^) for silica-coated AuNPs shows a small heat transfer enhancement (increased by 4 %) compared with the bare AuNP (*h*_*pw*_, 105 MW m^-2^K^-1^). However, the calculation of the PA generation (Figure 3C) does not show PA amplification despite the high silica-water ITC. Moreover, when the silica shell thickness is larger than 5.5 nm, even considering the enhanced silica-water heat transfer, 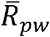 of silica coating will still be larger than *R*_*pw*_ due to the heat resistance in the thick silica shell (Figure S7). Also, for our experiment conditions (15 nm AuNP with 4 nm silica coating), considering *e-ph* coupling on the gold-silica interface alone does not explain the PA enhancement either (Figure 3C and D). Therefore, the combination of two interface heat transfer effects contributes to the PA enhancement, specifically the *e-ph* coupling effect on the gold-silica interface and enhance interface thermal conductance on the silica-water interface.

## Discussion

In this work, we performed a systematic investigation of how the pulse laser duration and silica coating thickness impact the PA amplification of spherical AuNPs. We experimentally observed a narrow window of PA amplification for AuNPs coated with thin (< 5 nm) silica shells under ps laser pulse stimulation. Theoretical modeling suggested that this unique regime of PA amplification can be explained by a synergistic combination of 1) enhanced interface heat transfer due to electron-phonon coupling at the gold-silica interface and 2) enhanced interface thermal conductance across the silica-water interface due to hydrogen bonding at the silica surface.

We have summarized how our results compare to the literature for experimental studies of PA signal alteration from Au@SiO2 nanoparticles in Table S2. In our work, we found that the PA amplification of spherical AuNPs due to the enhanced interface heat transfer effects of silica coating was limited to ps pulse laser stimulation with the silica coating having no discernible effect on the PA signal generation under ns pulse laser stimulation. However, as shown in Table S2, PA amplification has been reported in the literature for gold nanorods under ns pulse laser stimulation, and at least one report indicates PA amplification for gold nanostars under ns laser stimulation. Even though differences in the surface area to volume ratios between the different AuNP shapes may alter the importance of interface heat transfer, there are three other effects noted in the literature that may contribute to PA response enhancement of AuNPs with a silica coating including: i) a reduction in the energy threshold for cavitation and a resulting increased contribution of cavitation effects to the PA signal,^24,48^ ii) a shift in the absorbance peak that increases the AuNP power generation,^49^ and iii) an increase in the photothermal stability of the particles which reduces degradation of the PA signal during repeated laser pulse applications.^50^ Experimentally discerning the impact of interface heat transfer on the PA amplification of non-spherical silica-coated AuNPs will likely require additional measures like those implemented in our study to mitigate these effects. We have also summarized the literature for theoretical studies of PA signal alteration from Au@SiO2 nanoparticles in Table S3. As shown there, the relevant modeling efforts have focused on spherical AuNPs and suggest that limited heat conduction through the silica should reduce the PA signal under ns laser pulse stimulation. Similarly, our modeling indicated a small decrease in the PA signal with silica coating thickness for spherical AuNPs under ns laser pulse stimulation (Figure 3F).

Although the experimental and theoretical results for ps laser stimulation both indicate that there is only significant PA enhancement with thin silica shells (< 5 nm), there are some quantitative differences between the simulation and experimental results of the 15 nm AuNP. For example, the simulations predict a monotonic decrease in the PA enhancement with silica shell thickness (for SiO2 > 0), while the experiments show an initial increase with silica shell thickness followed by a sharp drop in the PA amplification for silica shell thickness > 5 nm. There are several additional factors that may affect the PA enhancement that will require further investigations to resolve their impact. First, both the heat transfer and PA generation processes can be affected by the water content in the silica shell. The silica shell inevitably contains some water due to the synthesis process and porosity.^24,25^ In terms of heat transfer, the water content decreases the silica thermal conductivity. The thermal conductivity of porous silica can be much lower than that of the fused silica thermal conductivity which is used in our model. A previous study suggests that experimental data matches simulation better when a lower thermal conductivity for silica is used in the model to account for the effect of porosity on PA generation under ns laser.^35^ However, the porosity of silica synthesized by the Stöber method remains to be determined. For our case, the decrease of silica thermal conductivity will decrease the PA enhancement due to a higher equivalent interface thermal resistance. On the other hand, the water content in silica can also enhance PA generation because water confined to narrow silica nanopores will have enhanced thermal expansion. Garofalini *et al*. demonstrated that the thermal expansion of water confined in 3 nm pores leads to a ∼70% enhancement over that of the bulk counterpart.^51,52^ Second, the contribution of PA generation in gold and silica is not considered in our numerical model. Although more than 90% of the PA generation is expected to come from the surrounding medium for AuNPs exposed to ns laser excitation,^34,53^ the nanoparticle’s contribution to PA generation may be larger when shorter laser pulses are applied.^53^ For instance, more than 60% of the PA signal may be generated by the AuNP (20 nm) under a 15 ps laser pulse. However, this pulse duration dependence is only based on numerical simulation and has not been confirmed by experiments. When the acoustic wave was generated in solid, only a portion of acoustic energy will be transmitted into the surrounding medium, known as an acoustic impedance mismatch (Eq. 12, SI). For the gold and water interface, only 9% of the acoustic energy in gold can be transmitted to water. Therefore, it is questionable whether the acoustic energy generated in AuNPs can be effectively transmitted to the surrounding medium. The surrounding medium can significantly affect the nanoparticle acoustic vibration. For instance, Major *et al*. demonstrated that the gold nanowire acoustic damping is sensitive to the properties of a surrounding medium, especially the acoustic impedance of the medium.^54^ With a silica coating, the transmitted energy from gold to water increases from 9% to 20% due to the acoustic impedance between gold and water. This may also contribute to the photoacoustic enhancement observed in our experiments, but it is much smaller than the observed 4-fold enhancement.

Future work includes the assessment of PA amplification from different pore sizes of silica shells such as mesoporous and fused silica, the impact of surface ligands, as well as the exploration of potential biomedical applications such as neuron stimulation^55^ and blood-brain barrier modulation.^56^ Various reports suggest the presence of mechanosensitive channels such as TRPV4 and PIEZO on neurons, and the amplified photoacoustic signal may represent an exciting opportunity for these applications.

In summary, we systematically studied the effect of nanostructure and laser duration on photoacoustic signal generation. We observed that thin silica coating (< 5 nm) can enhance PA signal by up to 4-fold compared to bare AuNP under picosecond laser. Theoretical modeling suggests that two interface energy transport processes are responsible for the PA enhancement under picosecond laser, 1) the enhanced interface thermal conductance on the silica/water interface and 2) the energy transport channel between gold electron and silica phonon. This study is an important step towards a better understanding of PA signal generation from plasmonic nanoparticles by providing a clear mechanism to account for the PA enhancement by silica coating. Our findings are valuable for the nanoscience and PA communities to design highly efficient PA agents for biomedical and other applications such as neuromodulation.

## Supporting information

Figure S1-S7, Table S1-S4, Equation S1-S15

