## Supplementary material for "Significantly amplified photoacoustic effect for silica-coated gold nanoparticles by interface heat transfer mechanisms": Figure S1-S7, Table S1-S4, Equation S1-S15

**Significantly amplified photoacoustic effect for silica-coated gold nanoparticles by tuning laser pulse duration**

**Materials and Methods**

**Gold nanoparticles (AuNPs) synthesis**. 15 nm AuNP was prepared by following the Frens’ as previously reported.^1,2^ Briefly, 1 ml of HAuCl_4_∙3H_2_O solution (25 mM) was added into Erlenmeyer flask cleaned by aqua regia having 100 mL of ultrapure water and the solution was boiled under magnetic stirring. 1 mL of sodium citrate (112.2 mM) solution was quickly added into the boiling solution and vigorously kept stirring for 10 min under heat. After the reaction, the solution was cooled down to room temperature and was filled with ultrapure water to be 100 mL total. The particles were used for the synthesis of Au@SiO_2_ or 40 nm AuNPs.

**40 nm AuNPs were prepared by the method of a seed-mediated growth**.^3^ 15 nm AuNPs were used as the seeds. In brief, the 300 mL of ultrapure water in a clean round bottom flask was heated up 95℃ with a reflux condenser. 1.8 mL of sodium citrate solution (10 mg/mL) and 10 mL of 15 nm AuNP seed (2 nM) were added to the flask. Subsequently, 300 µL of HAuCl_4_∙3H_2_O (25 mM) was added dropwise to the solution and the solution kept stirring for 30 min. An additional 300 µL of HAuCl_4_∙3H_2_O was added to the reaction. The step of adding the citrate solution and gold ion solution was repeated 3 more times. The particle solution was then aged for 1 h under the heat. After cooling down to room temperature, the particles were kept at 4 ℃ for future use.

**Silica-coated gold nanoparticles (Au@SiO_2_) synthesis.** Silica-coated gold nanoparticles were followed by the procedures reported previously (Figure S1).^4^ Briefly, 100 mL of fresh 15 nm AuNP solution (2 nM) was placed on a round bottom flask. 1 mL of aminopropyl trimethoxy silane (APTMS, 1 mM) was dropwise added to the AuNP solution under vigorous magnetic stirring and the mixture was stirred for 2 h. Meanwhile, 0.53 wt% of sodium silicate solution was prepared and adjusted pH to be 10.2 by acidic cation exchanger. After adjusting pH, the ion exchanger was removed by a filter. 4 mL of the prepared sodium silicate solution was added to the reactor. The solution was kept stirring until the active silica coating reached 3 nm from DLS measurement. Generally, 48 h or 72 h reaction time is required to have 3 nm active silica coating. After forming 3 nm active silica coating, the particle solution was divided into 1.5 mL microcentrifuge tubes and centrifuged at 4000 g for 90 min. After removing the supernatant, the particles were collected in 20 mL the ultrapure water, which is 5 times concentrated nanoparticle suspension. For thicker silica shell growing by Stöber method, 17 mL of the concentrated particle suspension was added dropwise to the clean 250 mL round bottom flask with 68 mL of absolute ethanol. Subsequently, 680 µL of ammonium hydroxide solution (28-30%) was injected into the solution. After 5 min, 10 µL of TEOS was added and the solution is allowed to stand for 18 h under mild magnetic stirring. The additions of TEOS were repeated until the desired silica shell thickness is obtained. For the washing of particles, 0.5 mL of the particle solution was mixed with 1 mL of ultrapure water in 1.5 ml microcentrifuge tubes and then the mixture was centrifuged at 8,000 g for 30 min. Afterwards, the supernatant was removed, and the pellet was re-dispersed in ultrapure water. Subsequently, the solution was centrifugated at the same condition. The product in water (Au@SiO_2_) was kept at 4 ℃ for storage before further experiment.

**Characterization of AuNPs and Au@SiO_2_**. The UV-Vis spectra of AuNPs and Au@SiO_2_ was recorded by a microplate reader (Synergy 2, BioTek). The hydrodynamic diameter of particles was measured by Malvern ZetaSizer Nano ZS. The TEM images were taken using JEOL JEM-2010 microscope operated at 120 kV.

**Experimental setup for photoacoustic measurement.** Figure 1A and Figure S2 show our experimental setup for the photoacoustic signal detection. 3 cm length of dialysis tubing (MWCO: 8-10 kD, 6.4 mm diameter) connected with polyethylene tube on both sides was installed in a water bath. The laser beam with ~0.3 cm^2^ focus spot size was aligned on the middle of dialysis tubing. The Quantel Q-smart 450 Nd:YGA laser is used for the generation of ns pulsed laser (532 nm, full width half maximum (FWHM) = 6 ns, 2 Hz). The EKSPLA PL 2230 Nd:YGA is used for the generation of ps pulsed laser (532 m, FWHM = 28 ps, 2 Hz). The particle solution was prepared with matched 0.3 OD at 532 nm. Each particle solution was introduced to a polyethylene tube by a syringe pump with 4 mL/min rate. The photoacoustic signal generated from the particle suspension in the dialysis tubing was detected by an unfocused immerse transducer with 0.5 MHz central frequency made by Olympus Corporation (V318-SU). The transducer was positioned 20 mm above the dialysis tubing. From each sample, the photoacoustic signal was collected from 30 laser pulses. To compare the intensity of photoacoustic signal, we determine the signal gap between the highest peak and the lowest peak (peak-to-peak, p2p) in the waveform and calculate the average and standard deviation of the peak intensities of 30 pulses. A low-pass filter was applied to PA signal to filter out the noise.

PA enhancement (%) was calculated from PA signal intensity of sample ($P$), Average PA intensity of bare AuNP ($P_{Au}$), Average PA intensity of water ($P_{H_{2}O}$):

$$PA enhancement \left( \% \right)= \frac{P-P_{Au}}{P_{Au}- P_{H_{2}O}} \times100$$

(1)

**Finite element model.** The two-temperature heat transfer model was adapted from Chen *et al.* Briefly, parabolic two-step model (P2T) was used in this study and neglects the electron thermalization process in the gold nanoparticle since it occurs in the fs range. Chen *et al* has demonstrated that P2T is sufficient for pure metal when laser duration is much longer than electron relaxation time (in the range of 0.01~10s fs). The governing equations of the two-temperature heat transfer model is listed as following,

$$\rho_{Au}C_{e}\frac{\partial T_{e}}{\partial t}-\nabla\cdot k_{e}\nabla T_{e}=Q_{v}\left( t \right)-G_{ep}\left( T_{e}-T_{p} \right)$$

(2)

$$\rho_{Au}C_{p\_Au}\frac{\partial T_{p}}{\partial t}-\nabla\cdot k_{p}\nabla T_{p}=G_{ep}\left( T_{e}-T_{p} \right)$$

(3)

$$\rho_{silica}C_{s}\frac{\partial T_{s}}{\partial t}-\nabla\cdot k_{s}\nabla T_{s}=0$$

(4)

$$\rho_{m}C_{m}\frac{\partial T_{m}}{\partial t}-\nabla\cdot k_{m}\nabla T_{m}=0$$

(5)

$$\mathbf{n}\cdot k_{e}\nabla T_{e}=-h_{es}\left( T_{e}-T_{s} \right)$$

(6)

$$\mathbf{n}\cdot k_{p}\nabla T_{p}=-h_{ps}\left( T_{p}-T_{s} \right)$$

(7)

$$\mathbf{n}\cdot k_{s}\nabla T_{s}=-h_{sm}\left( T_{s}-T_{m} \right)$$

(8)

$$Q_{v}=\frac{C_{abs}F}{V_{NP}}\frac{3}{\mu\sqrt{2\pi}}e^{-\frac{{9\left( t-\mu\right)}^{2}}{2\mu^{2}}}$$

(9)

The PA generation and propagation is defined by equation 10,

$$\frac{1}{\rho_{m}{c_{s}}^{2}}\frac{\partial^{2}p_{w}}{\partial t^{2}}-\nabla\cdot\left( -\frac{1}{\rho_{m}}\nabla p_{w} \right)=\frac{\partial}{\partial t}\left( \beta\frac{\partial T_{m}}{\partial t} \right)$$

(10)

where cs is the speed of sound, β is thermal expansion coefficient of water and p_w_ is pressure in water.

The PA quantum yield is defined as

The p is pressure at the boundary of the water domain, z is the acoustic impedance of water, F is

$$\Phi_{PA}=\frac{E_{acoustic}}{E_{optical}}=\frac{4\pi{r_{m}}^{2}}{z}\frac{\int p^{2}dt}{FC_{abs}}$$

(11)

the laser fluence and C_abs_ is the absorption cross section of the gold core.

The transmitted energy can be roughly estimated by the following equation,

$$TE=1-\left( \frac{z_{2}-z_{1}}{z_{2}+z_{1}} \right)^{2}$$

(12)

where Z is acoustic impendence of the material. All parameters used in the numerical model are listed in Table S4.

The thermal resistance on an interface due to ITC can be calculated by

$$R=1/Ah$$

(13)

where A is the surface area of the interface and h is the ITC of the interface. Similarly, the thermal resistance in silica domain due to the conductance can be calculated by Eq.14

$$R_{s}=\frac{\left( 1/{r_{NP}} \right)-(1/{(r_{NP}+a_{s}}))}{4\pi k_{s}}$$

(14)

where *r_NP_* is the particle diameter and k_s_ is the thermal conductivity of silica.

The equivalent thermal resistance ($\bar{R}_{pw}$) is defined as

$$\bar{R}_{pw}=R_{ps}+R_{s}+R_{sw}$$

(15)

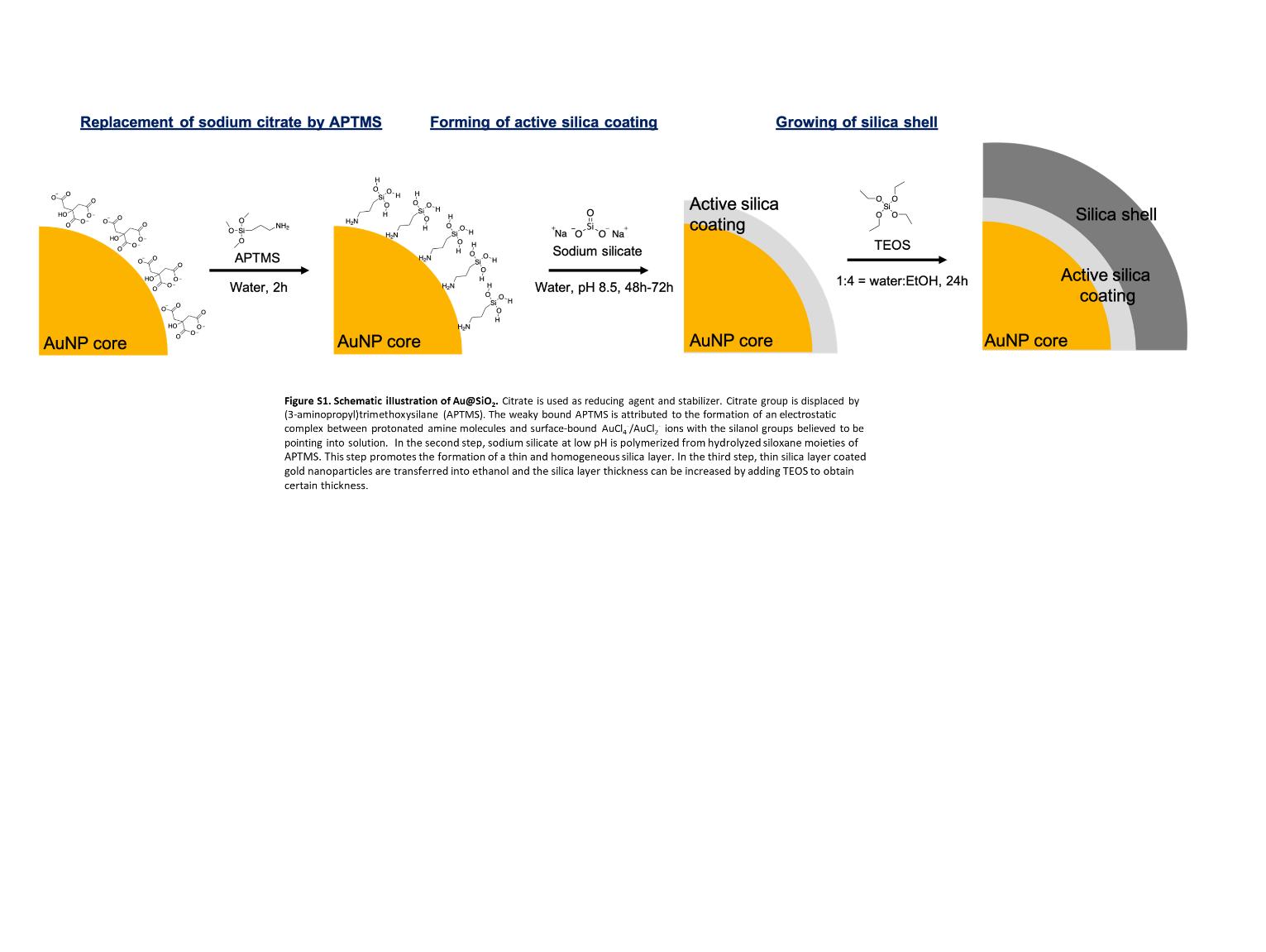
**Fig. S1. Schematic illustration for fabrication of Au@SiO_2_.** Citrate is used as a reducing agent and stabilizer. Citrate group is displaced by (3-aminopropyl)trimethoxysilane (APTMS). The weakly bound APTMS is attributed to the formation of an electrostatic complex between protonated amine molecules and surface-bound AuCl_4_^-^/AuCl_2_^-^ ions with the silanol groups believed to be pointing into solution. In the second step, sodium silicate at low pH is polymerized from hydrolyzed siloxane moieties of APTMS. This step promotes the formation of a thin and homogeneous silica layer. In the third step, thin silica layer coated gold nanoparticles are transferred into ethanol and the silica layer thickness can be increased by adding TEOS to obtain a certain thickness.

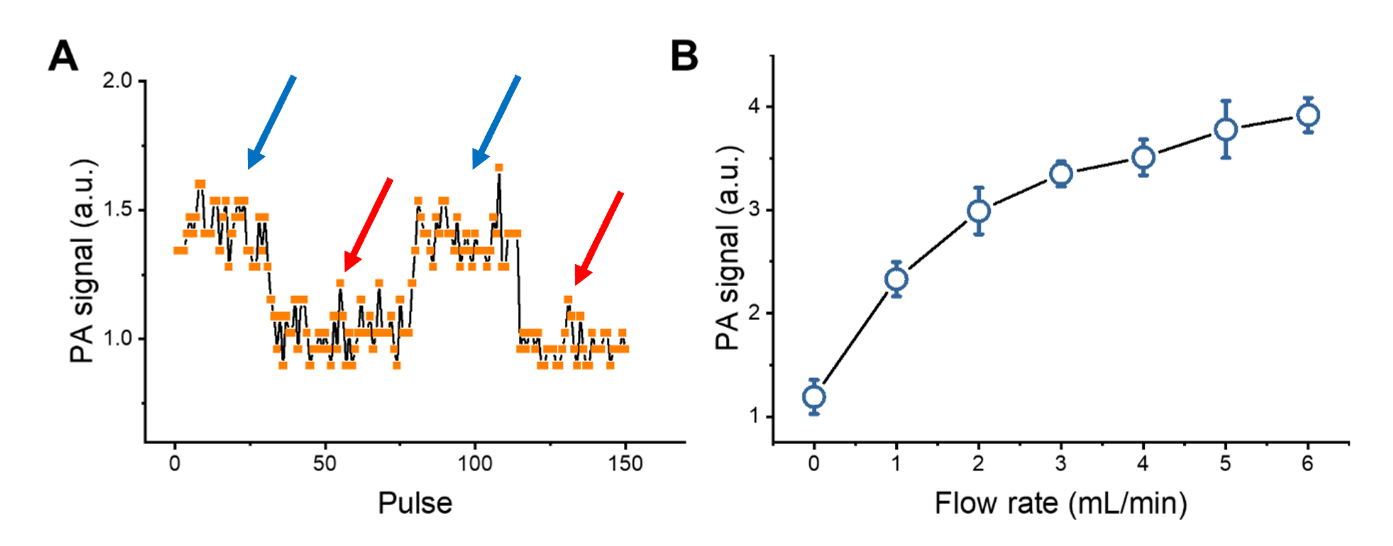

Fig. S2. The flow system for PA generation and detection. (A) The amplitude of PA signal of AuNPs with the flow (0.5 mL/min, blue arrows) and without flow (red arrows) under 19.7 mJ/cm^2^ ps laser irradiation. (B) The amplitude of PA signal of AuNPs generated from different flow rates with constant ps laser irradiation (19.7 mJ/cm^2^).

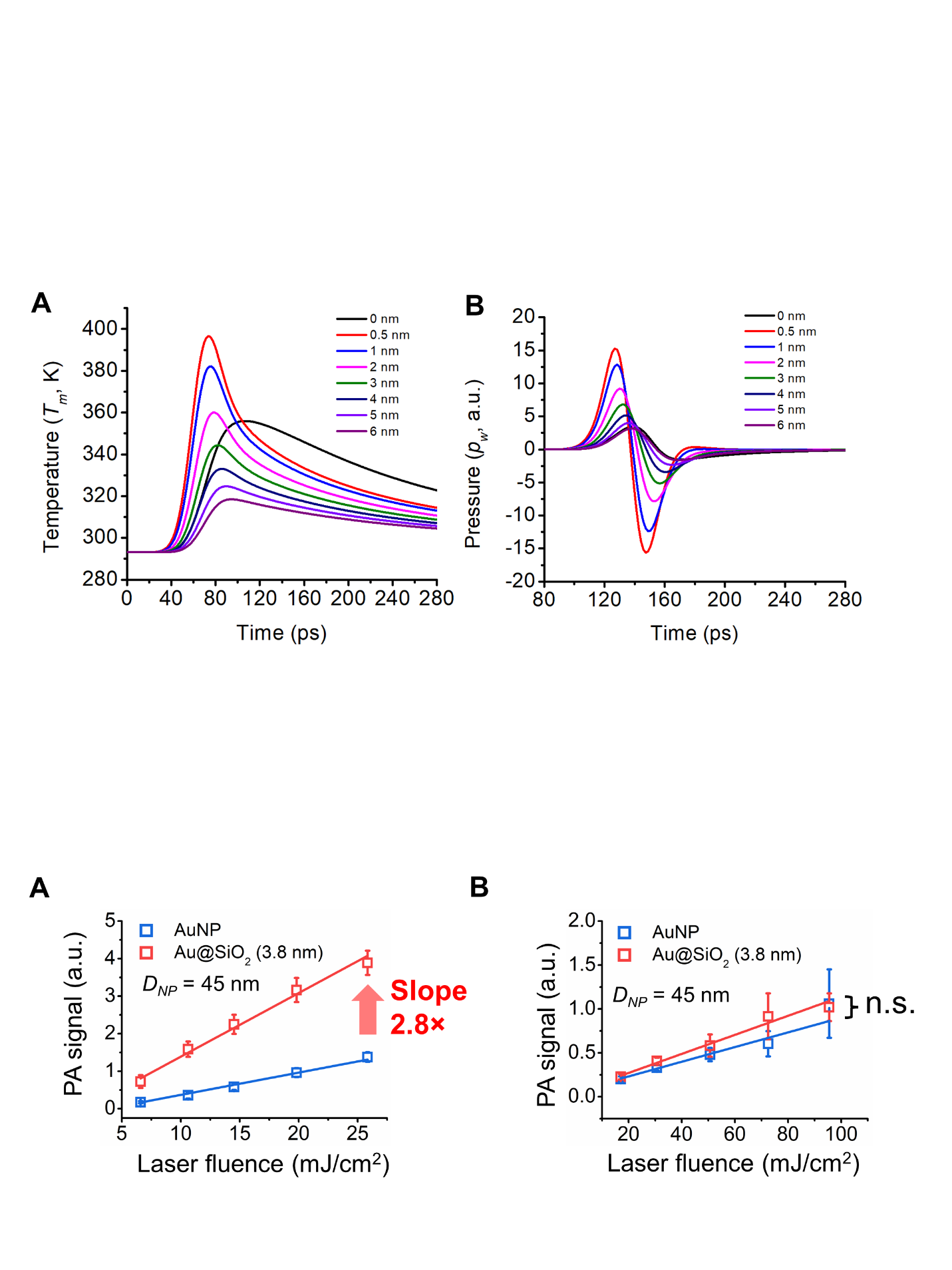

Fig. S3. Photoacoustic signal enhancement from silica-coated gold nanoparticles (45 nm Au core). The amplitude of photoacoustic signals of 15 nm bare AuNPs and Au@SiO_2_ induced by different laser fluences of picosecond laser (A) and nanosecond laser (B). n=30 measurements; the error bars show the standard deviation.

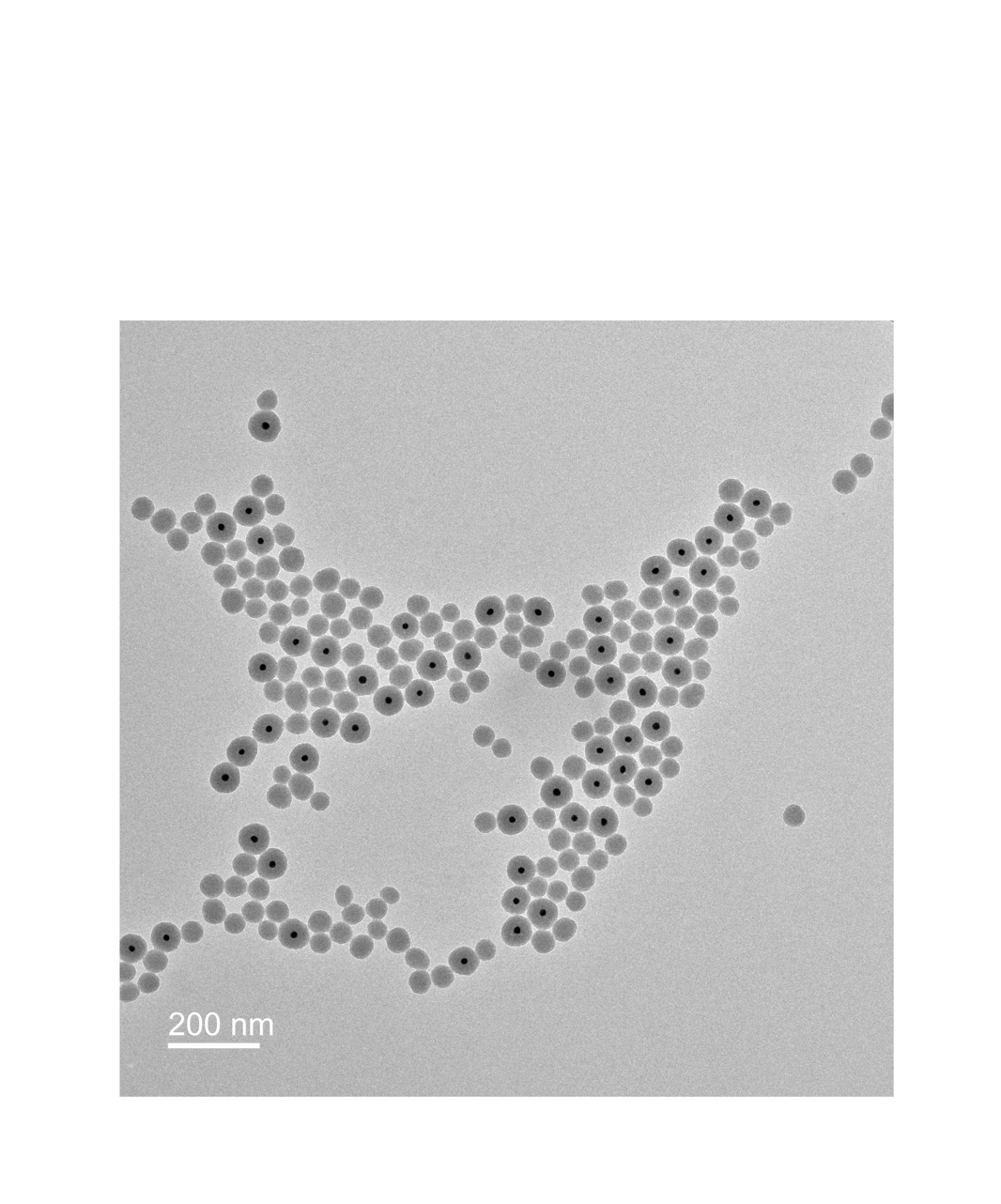
Fig. S4. TEM image of abundant silica nanoparticles after growing over 20 nm silica shell.

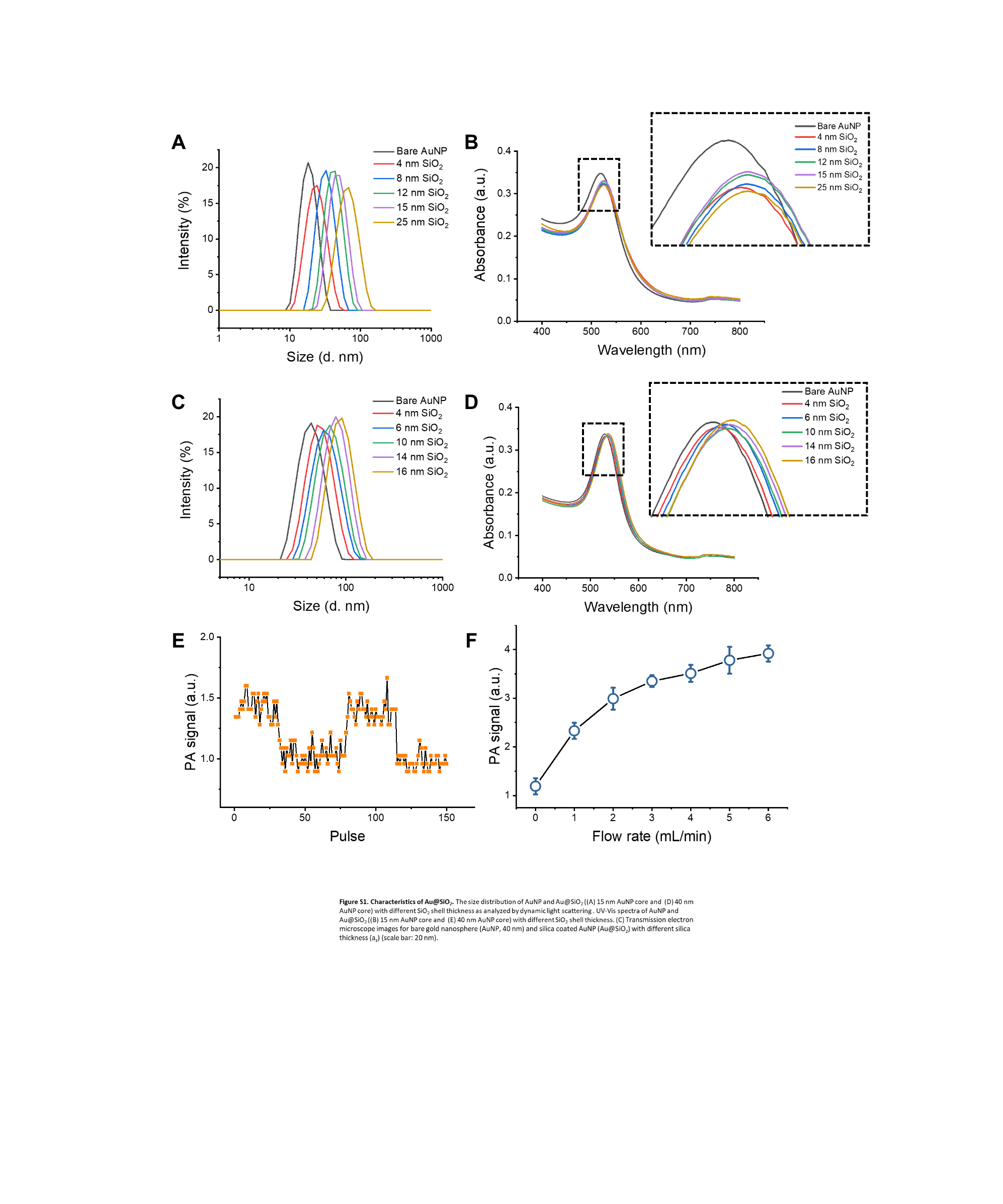
Fig. S5. Characterization of Au@SiO_2_ with 15 nm and 45 nm AuNP cores. The size distribution of AuNP and Au@SiO_2_ ((A) 15 nm AuNP core and (C) 45 nm AuNP core) with different SiO_2_ shell thickness as analyzed by dynamic light scattering (DLS). UV-Vis spectra of AuNP and Au@SiO_2_ ((B) 15 nm AuNP core and (D) 45 nm AuNP core) with different SiO_2_ shell thickness.

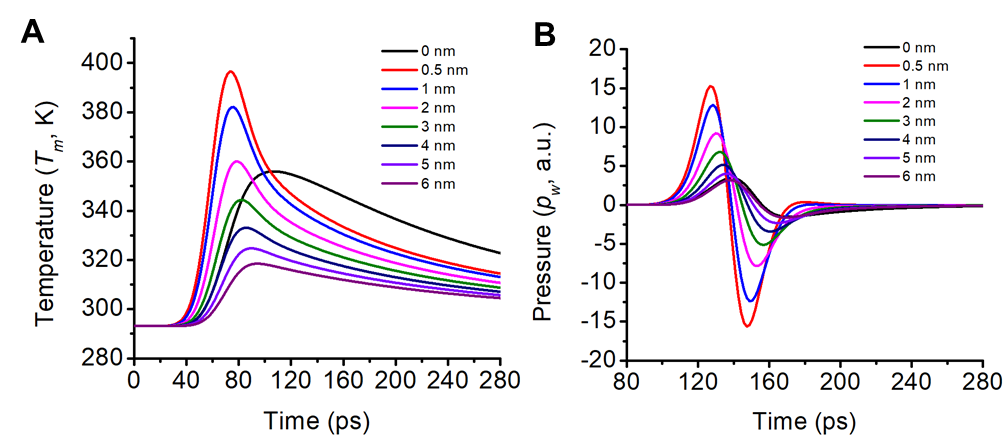
Fig. S6. (A) The temperature of the water on the surface of nanoparticles with different silica shell thicknesses. (B) The wavefunction of pressure at the boundary of water domain for nanoparticles with different silica shell thickness

Fig. S7.
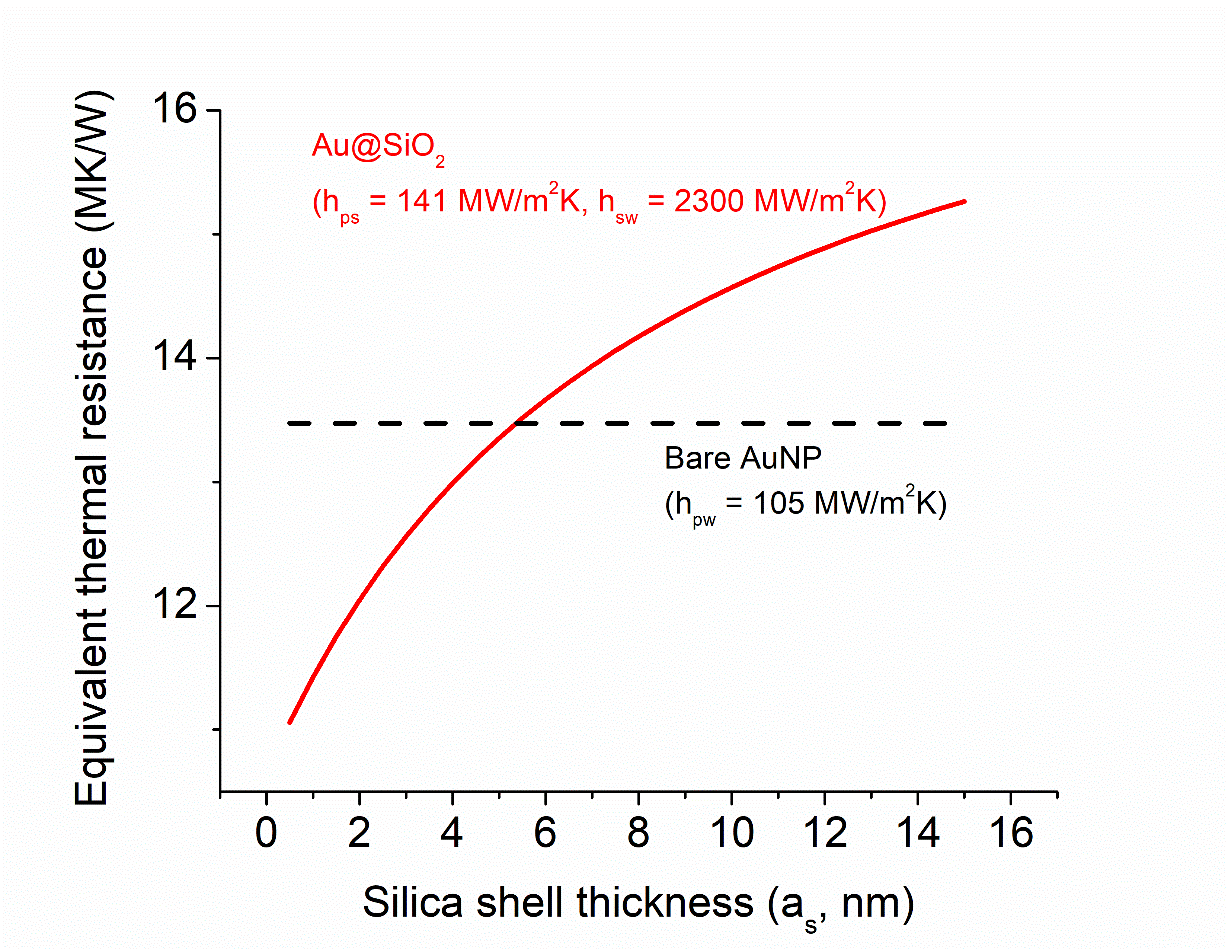
Equivalent thermal resistance for Au@SiO_2_ comparing with AuNP.

Table S1. Characterization of 15 nm and 45 nm core Au@SiO_2_.

| Core size  (nm) | SiO_2_ thickness determined by TEM  (nm) | DLS size  (d. nm) | PDI | Wavelength at max OD  (nm) |
| --- | --- | --- | --- | --- |
| 15 | 0 | 18.2 ± 0.3 | 0.02 | 518 |
|  | 3.9 | 24.0 ± 0.3 | 0.16 | 522 |
|  | 8.2 | 37.8 ± 0.9 | 0.11 | 523 |
|  | 12.1 | 43.6 ± 0.5 | 0.07 | 525 |
|  | 15.0 | 52.8 ± 0.1 | 0.04 | 525 |
|  | 25.4 | 76.7 ± 0.8 | 0.12 | 525 |
| 45 | 0 | 41.1 ± 0.1 | 0.09 | 530 |
|  | 4.9 | 50.1 ± 0.4 | 0.12 | 532 |
|  | 9.2 | 57.0 ± 0.4 | 0.12 | 534 |
|  | 10.5 | 64.3 ± 0.6 | 0.09 | 535 |
|  | 12.1 | 70.0 ± 0.4 | 0.07 | 536 |
|  | 16.8 | 80.4 ± 0.3 | 0.09 | 537 |

**Table S2. Comparison of experimentally reported PA signal alterations of Au@SiO_2_ nanoparticles upon pulsed laser irradiation.** Results from this work are included (bold text) for comparison.

| Shape of AuNP | Core Au diameter  [nm] | SiO_2_ thickness  [nm] | Pulse laser duration  [ns] | Laser power  [mJ/cm^2^] | Wavelength  [nm] | PA Amplification | Reported mechanism | Ref. |
| --- | --- | --- | --- | --- | --- | --- | --- | --- |
| Rod | 6 × 20 | 20 | 5 | 4 | 802 | 300% | Interface heat transfer | [5] |
| Sphere | 26 | 38 | 7 | 5 | 534 | 300% | Interface heat transfer | [6] |
| Sphere | 12.5 | 25 | 5 | 3.5 | 532 | - 25% | Silica heat conduction | [7] |
| Rod | 10x100 | 14 | 5-7 | 3.5 | 1064 | 15%-70% | Cavitation | [8] |
| Rod | 10x100 | 14  (hydrophobic coating) | 5-7 | 3-19 | 1064 | 475%-1258% | Cavitation | [8] |
| Rod | 15x24 | 20 | 4-6 | 40 | 680-970 | 400% | Peak shift | [9] |
| Star | 100 | 25 | 5-7 | 23 | 850 | 300% | Photothermal stability | [10] |
| **Sphere** | **15** | **4.6** | **0.028** | **6.6 – 25.8** | **532** | **400%** | **Interface heat transfer** | **This work** |
| **Sphere** | **15** | **4.6** | **6** | **17.0 –95.5** | **532** | **10%** | **Interface heat transfer** | **This work** |
| **Sphere** | **45** | **4.9** | **0.028** | **6.6 – 25.8** | **532** | **300%** | **Interface heat transfer** | **This work** |
| **Sphere** | **45** | **3.6** | **6** | **17.0 –95.5** | **532** | **50%** | **Interface heat transfer** | **This work** |

**Table S3. Comparison of PA signal alteration in simulation models for Au@SiO_2_ nanoparticles upon pulsed laser irradiation.**

| Shape of AuNP | Type of study | Core Au diameter  [nm] | SiO_2_ thickness  [nm] | Pulse laser duration  [ns] | Laser power  [mJ/cm^2^] | Wavelength  [nm] | Amplitude of PA signal | Ref. |
| --- | --- | --- | --- | --- | --- | --- | --- | --- |
| Sphere | Simulation | 5 - 20 | 10 - 50 | 5 | 1 | 532 | -25% | [7] |
| Sphere | Simulation | 30 | 5 - 30 | 10 | 2.7 | 532 | -10% | [11] |

**Table S4. Parameters used in numerical model**

| Symbol | Parameter | Value | Unit | Ref. |
| --- | --- | --- | --- | --- |
| $\rho_{Au}$ | Gold density | 19300 | Kg/m^3^ | [7] |
| $\rho_{silica}$ | Silica density | 2200 | Kg/m^3^ | [7] |
| $\rho_{m}$ | Water density | 1000 | Kg/m^3^ | [7] |
| $C_{e}$ | Specific heat of gold electrons | VSAP calculated | J/kg∙K | [12] |
| $C_{p\_Au}$ | Specific heat of gold phonons | 129 | J/kg∙K | [7] |
| $C_{s}$ | Specific heat of silica | 740 | J/kg∙K | [7] |
| $C_{m}$ | Specific heat of water | 4128 | J/kg∙K | [7] |
| $k_{e}$ | Thermal conductivity of gold electrons | VSAP calculated | W/m∙K | [12] |
| $k_{p}$ | Thermal conductivity of gold phonons | 2 | W/m∙K | [12] |
| $k_{s}$ | Thermal conductivity of silica | 1.38 | W/m∙K | [7] |
| $k_{m}$ | Thermal conductivity of water | 0.598 | W/m∙K | [7] |
| $h_{es}$ | Interface thermal conductance of gold electron-silica phonon coupling | 96.1+0.18T_e_ | MW/m^2^∙K | 12] |
| $h_{sw}$ (low) | Interface thermal conductance of silica-water interface | 150 | MW/m^2^∙K | [7] |
| $h_{sw}$ (enhanced) | Interface thermal conductance of silica-water interface (enhanced) | 2300 | MW/m^2^∙K | [13] |
| $h_{ps}$ | Interface thermal conductance of gold-silica interface | 141 | MW/m^2^∙K | [7] |
| $h_{pm}$ | Interface thermal conductance of gold-water interface | 105 | MW/m^2^∙K | [14] |
| $G_{ep}$ | Electron-phonon coupling factor | VSAP calculated | MW/m^3^∙K | [12] |
| $C_{abs}$ | Absorption cross section area | 158.5 | nm^2^ | [12] |
| $FWHM$ | Full width half maximum duration of laser pulse | 6 ns/28 ps | ns/ps |  |
| c_s_ | Speed of sound in water | 1500 | m/s | [7] |
| β | Thermal expansion coefficient | 1.6e-5+9.6e-6∙(T-273.15) | 1/K | [15] |
